# Collective effects of common SNPs and improved risk prediction in lung cancer

**DOI:** 10.1101/106864

**Authors:** Xiaoyun Lei, Dejian Yuan, Zuobin Zhu, Shi Huang

**Author notes:** These authors contributed equally to this work (Xiaoyun Lei, Dejian Yuan).

## Abstract

Lung cancer is the leading cause of cancer deaths in both men and women in the US. While most sporadic lung cancer cases are related to environmental factors such as smoking, genetic susceptibility may also play an important role and a number of lung cancer associated single nucleotide polymorphisms (SNPs) have been identified although many remain to be found. The collective effects of genome wide minor alleles of common SNPs, or the minor allele content (MAC) in an individual, have been linked with quantitative variations of complex traits and diseases. Here we studied MAC in lung cancer using previously published SNP datasets and found higher MAC in cases relative to matched controls. A set of 25883 SNPs with MA (MAF < 0.5) more common in cases (*P* < 0.1) was found to have the best predictive accuracy. A weighted risk score calculated by using this set can predict 2.6% of lung cancer cases (100% specificity). These results identify a novel genetic risk element or higher MAC in lung cancer susceptibility and provide a useful genetic method to identify a small fraction of lung cancer cases.

## Introduction

Lung cancer is the leading cause of cancer death in both men and women in the U.S and an estimated 158,040 Americans are expected to die from lung cancer in 2015, accounting for approximately 27 percent of all cancer deaths^1, 2^. The most common environmental risk factor for sporadic lung cancer is smoking and radon^3^. However, there are large variations in an individual’s susceptibility to lung cancer and the heritability of lung cancer is estimated to be 8-14%^4, 5^. Only a fraction of smokers (~15%) will develop lung cancer in their lifetime, and non-smokers also can develop lung cancers^6^. A number of cancer genes such as K-ras, p53, Rb, EGFR, HER2-neu have been identified whose mutations contribute to lung cancers^7-12^.

Efforts to identify quantitative susceptibility loci in lung cancer have mostly involved genome wide association studies (GWAS) and identified a number of lung cancer risk SNPs (single nucleotide polymorphisms, SNPs)^13-17^. However, they account for very small fraction of lung cancer cases and their mechanisms of action remain largely unknown^18^. Rather than focusing on individual major risk alleles, we have in recent years developed a novel approach to study the collective effects of weak effect SNPs on complex diseases and traits. We have shown that the collective effects of genome wide collection of minor alleles (MAs) in an individual are linked with lower reproductive fitness in *C*.*elegans* and yeasts^19^ and risk for Parkinson’s disease^20^. These studies suggest that overall level of randomness or MA amounts may be expected to be higher in complex diseases relative to controls.

Known predictive models of lung cancers mostly use smoking status, radon exposure, and family history^21, 22^. These models cannot predict pre-birth risk or risk long before incidence. Researchers have also used a set of susceptibility loci to create a genetic risk score to better predict lung cancer risk^23-25^. But these predictions were generally poor and not meaningful for clinical use. These prediction models have calculated the area under the receptor-operator curve (AUC). But they generally did not consider or could not generate meaningful true positive rate (TPR) with 100% specificity (no false positives), a more useful measure in clinical applications. Here we studied the overall level of genome wide randomness in lung cancer cases relative to controls as measured by total MA amounts in an individual. We also attempted to identify a set of MAs that can predict lung cancer risks.

## Materials and Methods

### SNPs datasets

We downloaded from dbGaP two case control GWAS datasets, phs000093.p2.v2 (Prostate, Lung, Colon and Ovary Study Cancer Screening Trial, PLCO)^26^ and phs000336.p1.v1 (Cancer Prevention Study II Nutrition Cohort, CPS-II)^16^. These studies used for SNP genotyping IlluminaHumanHap550v3.0, Human610_Quadv1_B, Human1M-Duov3_B. Cases were admitted based on chest X ray examination. Controls were matched healthy individuals with similar age, sex ratio, and location. These data were further cleaned by removing genetic outliers using Principle Component Analysis (PCA) with the GCTA tool^27^ (Supplementary Table S1 showing PCA values and plots). Also, SNPs were filtered by removing those with >5% non-informative calls in the population, and those not following the Hardy-Weinberg equilibrium in either the case group or the control group (*P*<0.0001 chi square test), and those with MAF<10E-6. Only autosome SNPs were used. The description of cleaned up datasets are shown in Table 1.

**Table 1.** Samples used in the study.

| Description | phs000093.v2.p2 |  | phs000336.v1.p1 |  |
| --- | --- | --- | --- | --- |
|  | Cases | Controls | Cases | Controls |
| Participants | 753 | 766 | 689 | 384 |
| SNPs | 535313 | 535313 | 535313 | 535313 |
| Male(%) | 472(62.7%) | 442(57.7%) | 386(56.0%) | 300(78.1%) |
| Age | - | - | 64.06±5.44 | 64.83±5.54 |

### Statistical analysis

Minor allele frequency (MAF) refers to the frequency at which the second most common allele occurs in a given population. MAs were defined as those alleles with MAF < 0.5 in the control group. The minor allele content (MAC) of an individual is the number of MAs divided by the total number of SNPs examined. We used a custom script^28^ to calculate the MAC values of case and control groups. Difference in the average MAC value was compared by two-sided Student’s *t* test. Each MA was given a weighted risk score calculated by logistic regression test, which was equal to the coefficient of the logistic regression test. For heterozygous MAs, the weighted risk score was 0.5 x the coefficient. The total MA numbers of each individual were then converted to a total weighted risk score by summing up the coefficient of each MA by using a custom script (Supplementary data 2).

### Haplotype construction

Haplotype block estimation was phased using PLINK^29^ with pairwise LD calculated for SNPs within 200kb. Haplotype selection was performed as described previously^30^. A standard logistic regression was performed on all SNPs of each haplotype to obtain association significance with the disease. For each haplotype, a representative SNP with the best disease linkage was chosen for risk prediction analysis and all SNPs chosen must satisfy the minimal selection criterion of MAF <0.5.

### Risk prediction

We performed two types of cross-validation experiments. For the external cross-validation analysis, the phs000093.v2.p2 cohort was used as a training set, and testing was performed on the phs000396.v1.p1 dataset. Each experiment’s discriminatory capability was evaluated using the receiver operating characteristic (ROC) curve. We then calculated the area under the curve (AUC) and the true positive rate (TPR) using Prism 6. TPR is the proportion of cases who had a risk score higher than that of any control individual. The AUC quantifies the overall ability of the test to discriminate between cases and controls. A truly useless test (one no better at identifying true positives than flipping a coin) has an area of 0.5. A perfect test (one that has zero false positives and zero false negatives) has an area of 1.00. In order to obtain a best MA set for risk prediction, six models were constructed using logistic regression. Five of these models were based on MAF, and the remaining one using haplotypes. We then obtained AUC and TPR of each set in the testing dataset phs000336.v1.p1.

In the internal 5-fold cross-validation analysis, the phs00093.v2.p2 cohort was randomly partitioned into 5 subsamples. Of the 5 subsamples, a single subsample was retained as the validation data for testing the model, and the remaining 4 subsamples were used as training data. The cross-validation process was then repeated 5 times, with each of the K subsamples used exactly once as the validation data. The 5 results were averaged to produce a single estimation. 10-fold cross-validation is commonly used and the advantage of this method over repeated random sub-sampling is that all observations are used for both training and validation, and each observation is used for validation exactly once. In account of the sample size in this study, 5-fold cross-validation was performed. Since the external cross validation analysis above identified the best MA set as having MAF < 0.5, we only analyzed this set of MAs in internal cross validation analysis. This set has 25,883 SNPs. As negative controls, we also used the whole set of non-selected 535,313 SNPs or 25,883 randomly selected SNPs for risk prediction.

### Pathway enrichment analysis

We used ANNOVAR^31^ to annotate the genes associated with the set of risk SNPs identified by the above analysis. We used DAVID^32^ to check the pathways associated with these genes in the KEGG (Kyoto Encyclopedia of Genes and Genomes)^33^. The enriched pathways in the risk SNPs set were compared by chi square test with those in the original dataset and a group of SNPs chosen randomly.

## Results

### Enrichment of minor alleles in lung cancer cases

We used two previously published GWAS datasets of lung cancer case and control cohorts for our studies. The cleaned datasets after removing genetic outliers were described in Table 1 (see Supplementary Table S1 for PCA plots). We merged these two datasets and then randomly divided the merged control dataset into two sets with 575 samples each. We used one of the control datasets for identifying minor allele status, and then calculated the MAC value of each individual in both case dataset and the two control datasets. We tested two sets of SNPs, one with MAF < 0.5 and the other with MAF < 0.5 but > 0.15. For SNPs set with MAF < 0.5, the average MAC value of control dataset 1 was lower than that of control dataset 2 but more significantly lower than that of cases (Table 2). For SNPs set with MAF > 0.15 and < 0.5, the average MAC value of control dataset 1 was not significantly different from that of control dataset 2 but significantly lower than that of cases (Table 2). These results indicate minor allele enrichment in lung cancer cases.

**Table 2.**
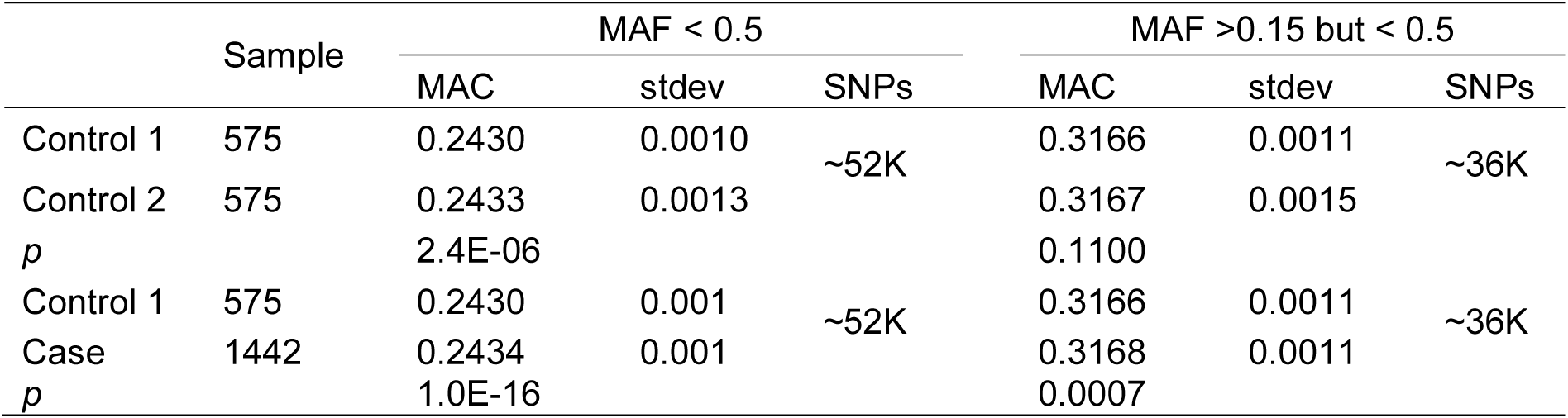
MAC value comparisons.

**Table 3.** Genes enriched in the lung cancer specific set of SNPs.

|  | Pathways |  |
| --- | --- | --- |
|  | ECM-receptor interaction | Endocytosis |
| Genes in prediction model | COL1A2 | ARFGAP2 |
|  | COL2A1 | CSF1R |
|  | COL11A2 | DAB2 |
|  | HMMR | FGFR2 |
|  | ITGA11 | LDLR |
|  | LAMA1 | PLD2 |
|  | LAMB1 | PSD |
|  | LAMB4 | PSD3 |
|  | TNC | RABEP1 |
|  | TNR |  |

We also studied the two downloaded datasets independently. For the dataset phs000093.v2.p2, we identified minor allele status by using the control group and calculated the MAC value of each case and control as well as the average MAC values of cases and controls. The results showed higher MAC values in cases than controls (*P* = 2.56E-07, Figure 1A and Supplementary Table S2). To confirm this result, we studied a second independent dataset phs000336.v1.p1, and obtained similar results (*P* = 1.46E-18, Figure 1B and Supplementary Table S2). Since case male ratio is different from it in control in second dataset phs000336.v1.p1, We compared male and female controls or cases and did not find significant differences in average MAC values between the sexes (Supplementary Table S3).

**Fig. 1.**
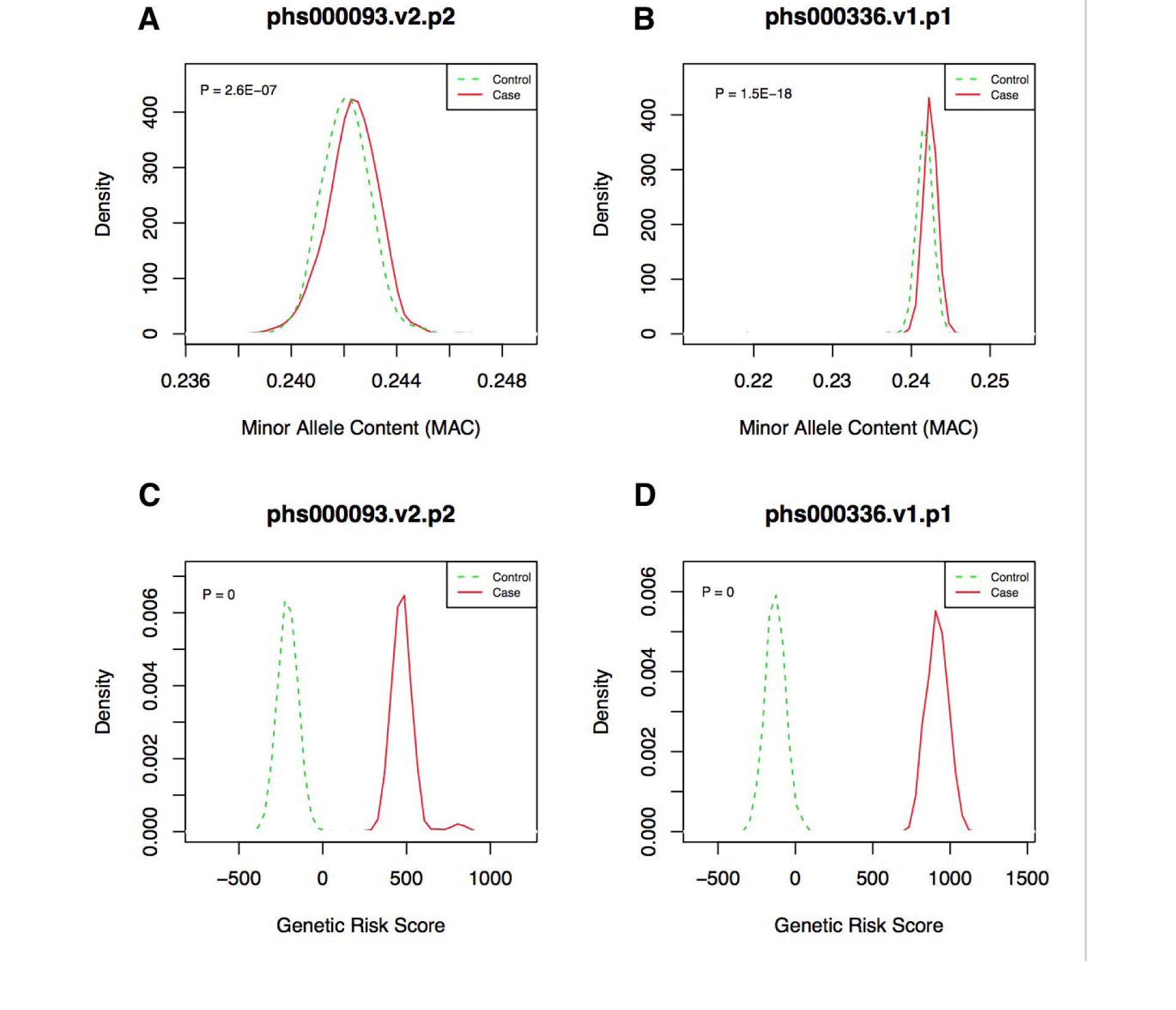
Distribution of MAC and genetic risk allele scores of MAs by case–control status. MAC: Minor allele content of SNPs with MAF < 0.5. Genetic risk score, the total risk score of all the MAs in an individual by adding the coefficient of logistic regression test of each MA.

Using logistic regression analysis, we obtained a weighted genetic risk score for each SNP. The total MA number of each individual was then converted into a total weighted risk score by summing up the coefficient of each MA (major alleles were not counted). The results showed clear separation of cases and controls in both datasets (Fig. 1C, 1D and Supplementary Table S2).

We next examined whether there is any relationship between MAF and the weighted genetic risk scores. We selected those SNPs with positive weighted risk score according to logistic regression and divided them into five groups based on their MAF values. For each group of SNPs, we calculated the total weighted risk score of each individual and obtained the average risk score of each SNP by dividing the total score by the number of SNPs examined. The result showed progressively higher risk scores for lower MAF values (Fig. 2), indicating that low frequency SNPs may be under more natural selection as a result of carrying higher risk of diseases.

**Fig. 2.**
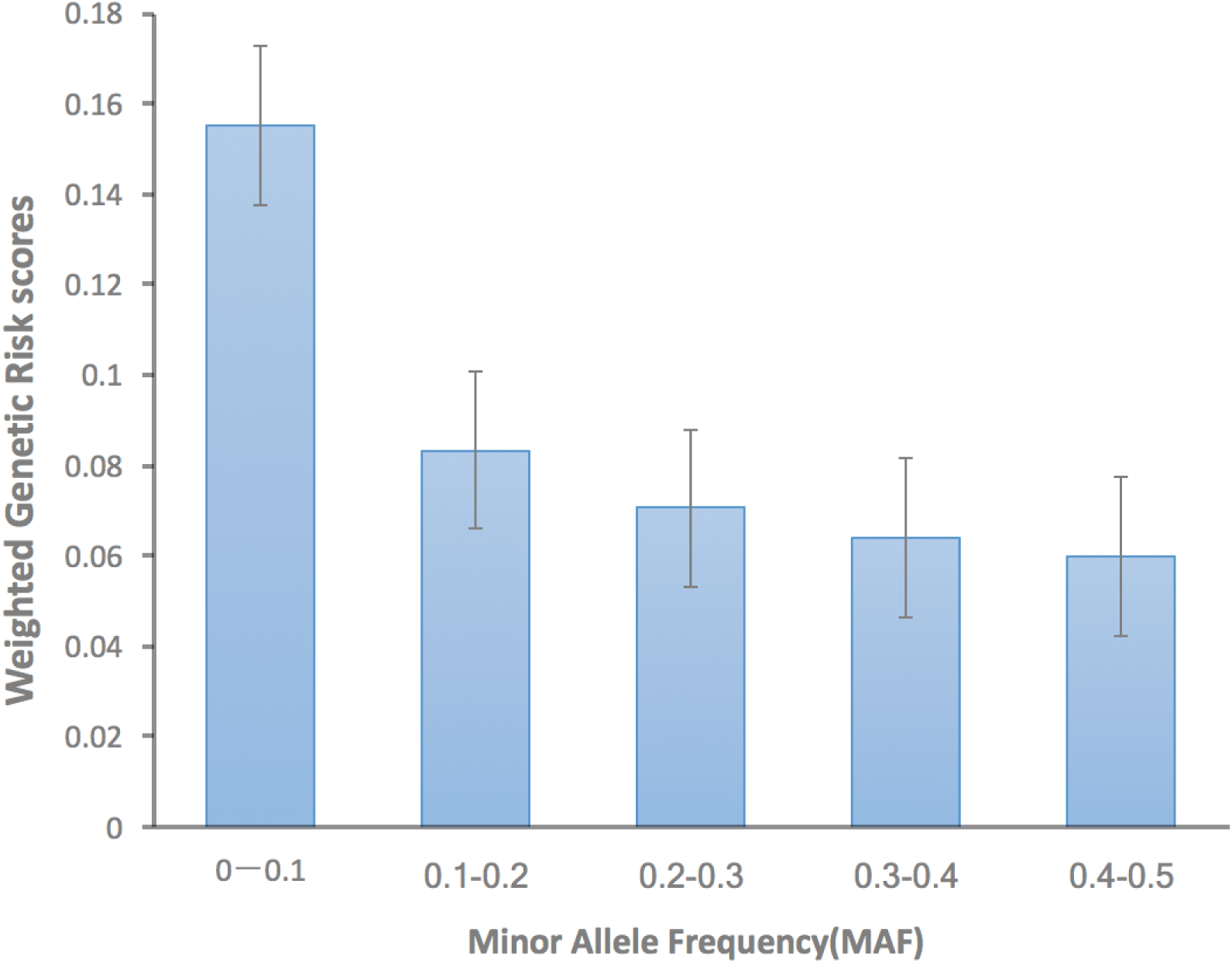
Relationships between MAF and lung cancer risk scores. Shown are the average weighted risk score for each category of MAs as classified by MAF.

### Risk prediction

We next aimed to obtain a specific set of MAs from a training dataset that could be used to predict lung cancer risk for an unrelated dataset (testing dataset). We chose the phs00093.p2.v2 dataset as the training dataset. In order to obtain a best MA set for risk prediction, six models were constructed using logistic regression. Five of these models were based on MAF, and the remaining one used haplotypes. We then used the receiver operator characteristic (ROC) curve and area under the curve (AUC) to examine the discriminatory capability of each set in the external cross validation analysis using the testing dataset phs000336.v1.p1. The SNP set showing the largest AUC as well as TPR was the one with MAF < 0.5 and each MAs’ linkage significance passing the threshold of *P* < 0.1 (Fig. 3, Supplementary Table S4). The AUC for this set is 0.5421 (95%CI, 0.5059-0.5782) and the TPR is 2.6% (95%CI, 1.555%-4.097%).

**Fig. 3.**
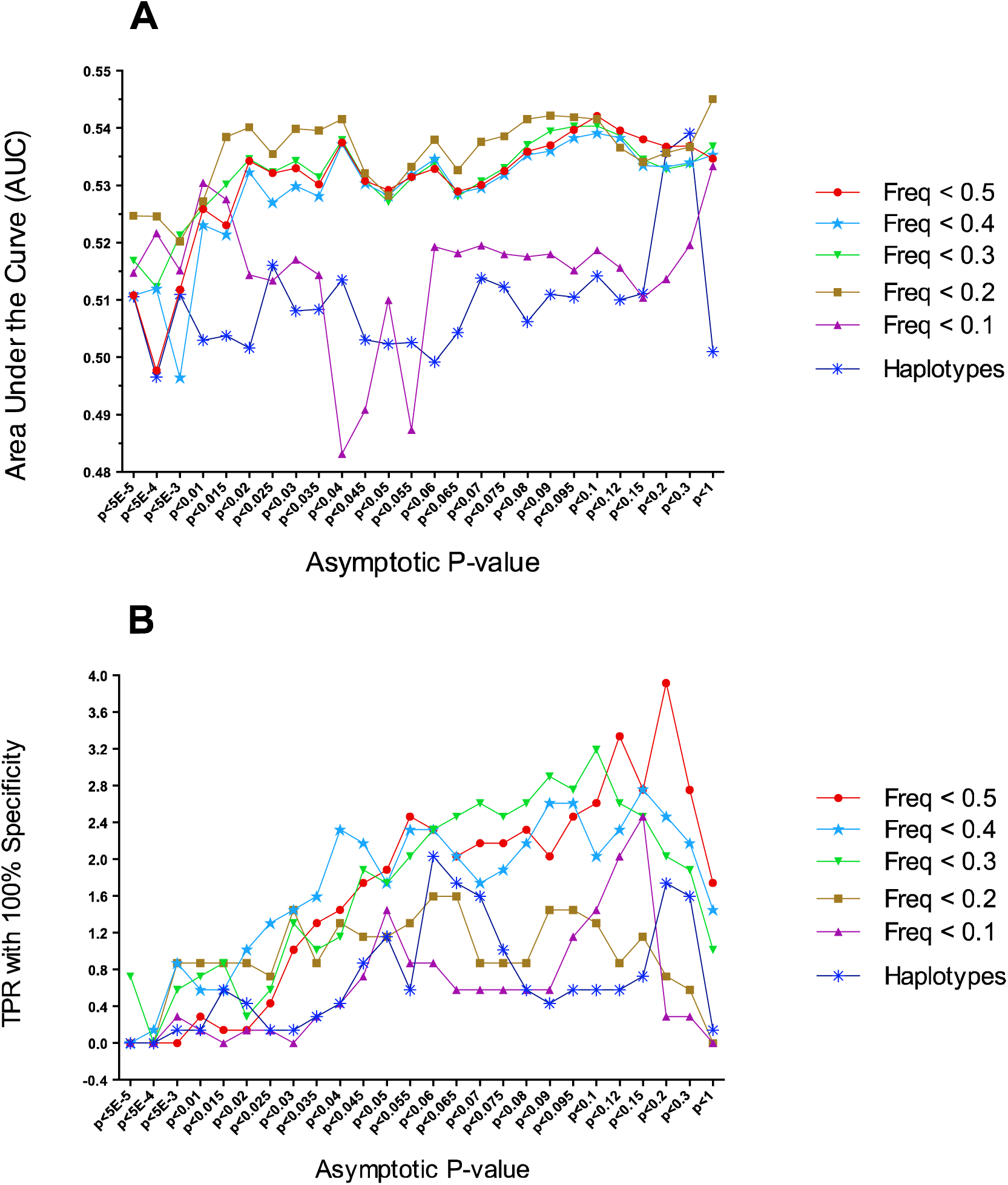
Discriminatory ability of different prediction models. SNPs were divided into 6 models based on MAF or haplotype. AUC (A) and TPR (B) were calculated using a training dataset and a test dataset. Each model was examined using MAs with different asymptotic P-value from the logistic regression test.

We further performed a 5 fold internal cross-validation analysis using the training dataset phs000093.v2.p2. Since the external cross validation analysis above identified the best set as having MAF < 0.5 with each MA passing the threshold of *P* < 0.1, we only analyzed this set in the internal cross validation analysis (see Supplementary Table S5 for risk scores of each SNP in the set). Using this set of 25883 SNPs a 5 fold internal cross-validation analysis, we obtained an average AUC of 0.5270 (95%CI, 0.5146-0.5394) and average TPR of 2.7% (95%CI, 1.818%-3.504%), which was similar to the above results of external cross validation analysis.

As negative controls, we performed external cross validation analysis by using the non-selected whole set of 535,313 SNPs, and found the TPR in this case to be only 1.7%. We also randomly selected 25883 SNPs and performed similar tests as we did for the above specifically selected set and found the average TPR to be only 1.08% (95%CI, 0.82%-1.33%).

### Pathway enrichment

Using ANNOVAR, we identified 354 genes in the set of 25883 risk SNPs as identified above. We then used DAVID to look for KEGG pathways associated with each of these genes. We identified two pathways that were enriched in the risk set relative to the original dataset, including extracellular matrix (ECM) with 10 genes, and endocytosis with 9 genes (Table 2).

We also analyzed the previously identified 37564 SNPs specific for Parkinson’s disease (PD)^20^. These SNPs were linked to 659 genes. We found only one pathway to be enriched in these genes, the Hedgehog signaling pathway with 6 genes. This pathway is known to be associated with PD^34-36^.

We next studied the lung cancer specificity of the 25,883 SNPs identified here by comparing it with the PD specific set of 37,564 SNPs as identified previously^20^. The non-selected whole set of SNPs (535,313 SNPs) in the lung cancer datasets here shared with the PD dataset (561,466 SNPs) 58.8% of SNPs. However, only 2.7% of SNPs were shared between the lung cancer specific set here and the PD specific set, indicating disease specificity (Supplementary Table S5).

### Discussion

Our results here suggest that having too many minor alleles of common SNPs at the genome wide level may be a novel genetic factor for lung cancer. We identified a lung cancer specific set of 25,883 SNPs that showed few overlaps with the previously identified SNPs specific for PD. Therefore, different diseases may be linked with different sets of MAs. While the number of SNPs involved in a disease may be quite large as indicated by this work here, much larger than expected from previous studies, it may not mean a lack of disease specificity in the collective effects of SNPs.

The result of higher MAC in lung cancer cases is a novel finding not expected by known works on human lung cancers. It confirms the previous result showing MAC association with lung cancer in a mouse lung cancer model^19^. Published lung cancer risk SNPs are relatively few in numbers. Therefore, even if these known risk alleles are mostly minor alleles, it may not predict that cases should have more genome wide MAs when a genome wide collection of ~500k SNPs are considered. If most MAs are not related to lung cancer except those few published lung cancer alleles, the average MAC of cases should not be significantly different from the controls.

Our study here further strengthens our intuitive hypothesis that a complex and ordered system must have an optimum limit on the level of randomness or entropy in its building parts or DNAs and the observation that human genetic diversities are presently at optimum level^20, 37-40^. While it may only take one or a few major effect errors to cause diseases, it would require the collective effects of a large number of minor effect errors in many different pathways to achieve a similar outcome. Cancer is known to be a disease of random mutations. Individuals with too many inherited random mutations or MAs may need fewer somatic mutations to pass the cancer threshold and hence have high susceptibility to cancer.

Negative selection by way of common diseases such as lung cancer may be one of the ways to maintain an optimum limit on genomic entropy. Although cancers tend to be late onset and hence well past the age of reproduction, which may be expected to have little selective effects on genes, it is easy to find ways for late onset common diseases to prevent accumulation of disease risk alleles in a population. For example, individuals with too many MAs may be already at a fitness disadvantage in many different traits including reproduction prior to disease onset at older age^19, 30, 41^.

Although AUC has been used in many studies for gauging performance of prediction model, some authors think that the AUC also has disadvantages^20, 25, 42, 43^. Our predictive model of lung cancer is comparable to previous results as indicated by AUC values^24^ and achieves a TPR of 2.6%. It has been shown that the prediction quality can be improved when there are a large number of SNPs^44^. Our prediction model for lung cancer also follows this trend with the prediction quality peaking at the case-association significance threshold of *P*<0.1. Models using small number of SNPs may be more susceptible to influence by random effects, while using too large number of SNPs may contain many irrelevant SNPs. A strong model may require a fine balance between high amounts of lung cancer linked SNPs and low amounts of irrelevant SNPs. Our best predictor model has a TPR value of 2.6% or could detect about 2.6% lung cancer patients as verified by both external and internal cross-validation experiment. It seems to be low but may still be meaningful. The value is higher than that of BRAF mutation with a mutation rate of 2.37% lung cancer cases^45, 46^.

The 25883 SNPs in our lung cancer prediction model were enriched in ECM-receptor interaction pathway and Endocytosis pathway. In contrast, randomly chosen SNPs of the same number did not have the same pathway enrichment. ECM-receptor interaction pathway includes extracellular matrix such as collagen, fibronectin and laminin, and transmembrane receptor such as integrin and proteoglycans. Most of these molecules are known to play roles in cancer^47-51^. Our results provide additional evidence for the role of these genes in lung cancer and may help understand their mechanisms of action in lung cancer.

## Acknowledgements

This work was supported by the National Natural Science Foundation of China (Grant No. 81171880) and the National Basic Research Program of China (Grant No. 2011CB51001).

The datasets used for the analyses described in this manuscript were obtained from dbGaP at http://www.ncbi.nlm.nih.gov/sites/entrez?Db=gap.The dbGaP accession numbers include phs000093.p2.v2 and phs000336.p1.v1. We thank PLCO Trial dbGaP GWAS Data, CPS-II dbGaP GWAS Data and the Contributing Investigator (phs000093.p2.v2, Neil Caporaso, Maria Teresa Landi, Lynn Goldin, etc; phs000336.p1.v1,Maria Teresa Landi, Neil E. Caporaso, Michael Thun, etc.). The content is solely the responsibility of the authors and does not necessarily represent the official views of the funding agencies.

## Supplementary Data

1. Supplementary Table S1. Principal component analysis (PCA).

2. Supplementary Table S2. The MAC and genetic risk scores of the individuals in the two case control datasets.

3. Supplementary Table S3. MAC values in male versus females.

4. Supplementary Table S4. Summary of AUC and TPR values.

5. Supplementary Table S5. MAs set for prediction in lung cancer and compared to PD

6. Supplementary materials. The script for calculating the risk score of each individual.

## References

1 Centers for Disease Control and Prevention. National Center for Health Statistics. CDC WONDER Online Database, compiled from Compressed Mortality File 1999-2012 Series 20 No. 2R, 2014.

2 American Cancer Society. Cancer facts and figures, 2015.

3 Alberg AJ, Samet JM. Epidemiology of lung cancer. Chest Journal 2003; 123: 21S–49S.

4 Czene K, Lichtenstein P, Hemminki K. Environmental and heritable causes of cancer among 9.6 million individuals in the Swedish family-cancer database. International Journal of Cancer 2002; 99: 260–266.

5 Hemminki K, Lönnstedt I, Vaittinen P, Lichtenstein P. Estimation of genetic and environmental components in colorectal and lung cancer and melanoma. Genetic epidemiology 2001; 20: 107–116.

6 Spitz MR, Wei Q, Dong Q, Amos CI, Wu X. Genetic susceptibility to lung cancer: the role of DNA damage and repair. Cancer epidemiology, biomarkers & prevention : a publication of the American Association for Cancer Research, cosponsored by the American Society of Preventive Oncology 2003; 12: 689–698.

7 Johnson L, Mercer K, Greenbaum D, Bronson RT, Crowley D, Tuveson DA et al. Somatic activation of the K-ras oncogene causes early onset lung cancer in mice. Nature 2001; 410: 1111–1116.

8 Takahashi T, Nau MM, Chiba I, Birrer MJ, Rosenberg RK. p53: a frequent target for genetic abnormalities in lung cancer. Science (New York, NY) 1989; 246: 491.

9 Iggo R, Bartek J, Lane D, Gatter K, Harris AL. Increased expression of mutant forms of p53 oncogene in primary lung cancer. The Lancet 1990; 335: 675–679.

10 Brabender J, Danenberg KD, Metzger R, Schneider PM, Park J, Salonga D et al. Epidermal growth factor receptor and HER2-neu mRNA expression in non-small cell lung cancer is correlated with survival. Clinical Cancer Research 2001; 7: 1850–1855.

11 Paez JG, Jänne PA, Lee JC, Tracy S, Greulich H, Gabriel S et al. EGFR mutations in lung cancer: correlation with clinical response to gefitinib therapy. Science (New York, NY) 2004; 304: 1497–1500.

12 Ding L, Getz G, Wheeler DA, Mardis ER, McLellan MD, Cibulskis K et al. Somatic mutations affect key pathways in lung adenocarcinoma. Nature 2008; 455: 1069–1075.

13 Amos CI, Wu X, Broderick P, Gorlov IP, Gu J, Eisen T et al. Genome-wide association scan of tag SNPs identifies a susceptibility locus for lung cancer at 15q25. 1. Nature genetics 2008; 40: 616–622.

14 Zhu Y, Hoffman A, Wu X, Zhang H, Zhang Y, Leaderer D et al. Correlating observed odds ratios from lung cancer case–control studies to SNP functional scores predicted by bioinformatic tools. Mutation Research/Fundamental and Molecular Mechanisms of Mutagenesis 2008; 639: 80–88.

15 Ryan BM, Robles AI, McClary AC, Haznadar M, Bowman ED, Pine SR et al. Identification of a functional SNP in the 3′ UTR of CXCR2 that is associated with reduced risk of lung cancer. Cancer research 2015; 75: 566–575.

16 Landi MT, Chatterjee N, Yu K, Goldin LR, Goldstein AM, Rotunno M et al. A genome-wide association study of lung cancer identifies a region of chromosome 5p15 associated with risk for adenocarcinoma. The American Journal of Human Genetics 2009; 85: 679–691.

17 Hu Z, Wu C, Shi Y, Guo H, Zhao X, Yin Z et al. A genome-wide association study identifies two new lung cancer susceptibility loci at 13q12. 12 and 22q12. 2 in Han Chinese. Nature genetics 2011; 43: 792–796.

18 Gibson G. Rare and common variants: twenty arguments. Nature Reviews Genetics 2012; 13: 135–145.

19 Yuan D, Zhu Z, Tan X, Liang J, Zeng C, Zhang J et al. Scoring the collective effects of SNPs: association of minor alleles with complex traits in model organisms. Science China Life Sciences 2014; 57: 876–888.

20 Zhu Z, Yuan D, Luo D, Lu X, Huang S. Enrichment of minor alleles of common SNPs and improved risk prediction for Parkinson’s disease. PloS one 2015; 10: e0133421.

21 Spitz MR, Hong WK, Amos CI, Wu X, Schabath MB, Dong Q et al. A risk model for prediction of lung cancer. Journal of the National Cancer Institute 2007; 99: 715–726.

22 Cassidy A, Myles JP, van Tongeren M, Page R, Liloglou T, Duffy S et al. The LLP risk model: an individual risk prediction model for lung cancer. British journal of cancer 2008; 98: 270–276.

23 Weissfeld JL, Lin Y, Lin H-M, Kurland BF, Wilson DO, Fuhrman CR et al. Lung cancer risk prediction using common SNPs located in GWAS-identified susceptibility regions. Journal of Thoracic Oncology 2015; 10: 1538–1545.

24 Li H, Yang L, Zhao X, Wang J, Qian J, Chen H et al. Prediction of lung cancer risk in a Chinese population using a multifactorial genetic model. BMC medical genetics 2012; 13: 118.

25 Jostins L, Barrett JC. Genetic risk prediction in complex disease. Human molecular genetics 2011; 20: R182–R188.

26 Caporaso N LM, Goldin L, et al. A Genome Wide Scan of Lung Cancer and Smoking, 2010. (https://www.ncbi.nlm.nih.gov/projects/gap/cgi-bin/study.cgi?study_id=phs000091.v2.p1)

27 Yang J, Lee SH, Goddard ME, Visscher PM. GCTA: a tool for genome-wide complex trait analysis. The American Journal of Human Genetics 2011; 88: 76–82.

28 health1987. dist: A tool to calculate nucleotide diversity 2016.( https://github.com/health1987/dist)

29 Purcell S, Neale B, Todd-Brown K, Thomas L, Ferreira MA, Bender D et al. PLINK: a tool set for whole-genome association and population-based linkage analyses. The American Journal of Human Genetics 2007; 81: 559–575.

30 Kang J, Kugathasan S, Georges M, Zhao H, Cho JH. Improved risk prediction for Crohn’s disease with a multi-locus approach. Human molecular genetics 2011; 20: 2435–2442.

31 Wang K, Li M, Hakonarson H. ANNOVAR: functional annotation of genetic variants from high-throughput sequencing data. Nucleic acids research 2010; 38: e164–e164.

32 Huang DW, Sherman BT, Lempicki RA. Systematic and integrative analysis of large gene lists using DAVID bioinformatics resources. Nature protocols 2009; 4: 44–57.

33 Kanehisa M, Goto S. KEGG: kyoto encyclopedia of genes and genomes. Nucleic acids research 2000; 28: 27–30.

34 Frank-Kamenetsky M, Zhang XM, Bottega S, Guicherit O, Wichterle H, Dudek H et al. Small-molecule modulators of Hedgehog signaling: identification and characterization of Smoothened agonists and antagonists. Journal of biology 2002; 1: 10.

35 Wu X, Walker J, Zhang J, Ding S, Schultz PG. Purmorphamine induces osteogenesis by activation of the hedgehog signaling pathway. Chemistry & biology 2004; 11: 1229–1238.

36 Bak M, Hansen C, Tommerup N, Larsen LA. The Hedgehog signaling pathway–implications for drug targets in cancer and neurodegenerative disorders. Pharmacogenomics 2003; 4: 411–429.

37 Huang S. The genetic equidistance result of molecular evolution is independent of mutation rates. Journal of computer science and systems biology 2008; 1: 92.

38 Huang S. Inverse relationship between genetic diversity and epigenetic complexity 2009. (Preprint available, Nature Precedings; http://precedings.nature.com/documents/1751/version/2)

39 Huang S. New thoughts on an old riddle: What determines genetic diversity within and between species? Genomics 2016; 108: 3–10.

40 Yuan D, Lei X, Gui Y, Zhu Z, Wang D, Yu J et al. Modern human origins: multiregional evolution of autosomes and East Asia origin of Y and mtDNA. bioRxiv 2017: 101410.

41 Meigs JB, Shrader P, Sullivan LM, McAteer JB, Fox CS, Dupuis J et al. Genotype score in addition to common risk factors for prediction of type 2 diabetes. New England Journal of Medicine 2008; 359: 2208–2219.

42 Pepe MS, Janes HE. Gauging the performance of SNPs, biomarkers, and clinical factors for predicting risk of breast cancer. Oxford University Press, 2008.

43 Hand DJ. Evaluating diagnostic tests: the area under the ROC curve and the balance of errors. Statistics in Medicine 2010; 29: 1502–1510.

44 Evans DM, Visscher PM, Wray NR. Harnessing the information contained within genome-wide association studies to improve individual prediction of complex disease risk. Human molecular genetics 2009; 18: 3525–3531.

45 Brose MS, Volpe P, Feldman M, Kumar M, Rishi I, Gerrero R et al. BRAF and RAS mutations in human lung cancer and melanoma. Cancer research 2002; 62: 6997–7000.

46 Molina JR, Yang P, Cassivi SD, Schild SE, Adjei AA. Non-small cell lung cancer: epidemiology, risk factors, treatment, and survivorship. Mayo Clinic Proceedings, vol. 83. Elsevier, 2008, pp 584–594.

47 Desgrosellier JS, Cheresh DA. Integrins in cancer: biological implications and therapeutic opportunities. Nature Reviews Cancer 2010; 10: 9–22.

48 Fang M, Yuan J, Peng C, Li Y. Collagen as a double-edged sword in tumor progression. Tumor Biology 2014; 35: 2871–2882.

49 Cox TR, Erler JT. Remodeling and homeostasis of the extracellular matrix: implications for fibrotic diseases and cancer. Disease models & mechanisms 2011; 4: 165–178.

50 Mosesson Y, Mills GB, Yarden Y. Derailed endocytosis: an emerging feature of cancer. Nature Reviews Cancer 2008; 8: 835–850.

51 Mellman I, Yarden Y. Endocytosis and cancer. Cold Spring Harbor perspectives in biology 2013; 5: a016949.

